# On forensic likelihood ratios from low-coverage sequencing

**DOI:** 10.1101/2024.05.24.595821

**Authors:** Feriel Ouerghi, Dan E. Krane, Michael D. Edge

**Affiliations:** Department of Quantitative and Computational Biology, University of Southern California; Department of Biological Sciences, Wright State University

## Abstract

Advances in sequencing technology are allowing forensic scientists to access genetic information from increasingly challenging samples. A recently published computational approach, `IBDGem`, analyzes sequencing reads, including from low-coverage samples, in order to arrive at likelihood ratios for human identification. Here, we show that likelihood ratios produced by `IBDGem` are best interpreted as testing a null hypothesis different from the traditional one used in a forensic genetics context. In particular, rather than testing the hypothesis that the sample comes from a person unrelated to the person of interest, `IBDGem` tests the hypothesis that the sample comes from an individual who is included in the reference database used to run the method. This null hypothesis is not generally of forensic interest, because the defense hypothesis is not typically that the evidence comes from an individual included in a reference database. Moreover, the computed likelihood ratios can be much larger than likelihood ratios computed for the standard forensic null hypothesis, often by many orders of magnitude, thus potentially creating an impression of stronger evidence for identity than is warranted. We lay out this result and illustrate it with examples, giving suggestions for directions that might lead to likelihood ratios that test the typical defense hypothesis.

## Introduction

One crucial goal of forensic genetics is identity testing, in which a biological sample can be tested to see whether the genetic data extracted from it could have originated from a person of interest (Balding & Donnelly, 1995; Gill et al., 1985; Jobling & Gill, 2004). The results are often expressed as a likelihood ratio comparing two hypotheses, with the prosecution’s hypothesis in the numerator and a defense hypothesis in the denominator (Balding & Steele, 2015). Most typically, in the numerator, the hypothesis is that the biological sample comes from a person of interest (e.g. a suspect or a victim), and in the denominator, the hypothesis is that the biological sample comes from a randomly chosen person from a population of alternative sources who is unrelated to the person of interest. We refer below to likelihood ratios of this form as “standard” forensic likelihood ratios. A long forensic and judicial tradition has built up in the interpretation of these likelihood ratios. Historically, such tests are most often performed with short tandem repeat (STR) markers, also called microsatellites (Butler, 2005).

In the last 30 years, genotyping technologies have advanced rapidly, and it is now possible to extract genetic information from samples that would have been intractable previously (Haddrill, 2021). Alongside new ways of generating data from challenging samples, new methods of analysis have appeared. For example, probabilistic genotyping systems (PGS) are now frequently used to analyze STR data from low-template and/or mixed samples (Coble & Bright, 2019).

For very challenging samples, it may be possible to generate some sequencing reads even if PCR amplification, which is necessary for STR typing, fails. In such a scenario, the sequencing depth may be too low to allow accurate genotype calls (Nielsen et al., 2011) thereby precluding the use of many existing methods and requiring special tools.

Recently, a handful of methods have appeared to address the possibility of computing forensic likelihood ratios from sequencing data. Mostad & colleagues (2023) extended the `Familias` package (Kling et al., 2014) to pedigree inference with low-coverage sequencing reads, assuming linkage equilibrium (but including linkage) among included sites. Andersen & colleagues (2025) presented a method for identity inference from unlinked markers that can account for genotyping errors from next-generations sequencing data, though it requires called genotypes. Nguyen and colleagues’ (2023) `IBDGem` is another such method. Like Mostad & colleagues’ extension of `Familias, IBDGem` works specifically with sequencing reads (from an unmixed sample) rather than called genotypes, an important extension that allows the analysis of samples in which only very low-depth sequencing data are attainable. Further, with its “LD mode,” `IBDGem` aims to accommodate sites in linkage disequilibrium (LD), unlike existing methods. `IBDGem` was developed with forensics in mind as a primary application area. Nguyen and colleagues developed a likelihood ratio statistic and evaluated its ability to distinguish identity from non-identity comparisons both within their reference data, with simulated lower coverage, and in real genotypes recovered from rootless hair and saliva. However, as we show here, the likelihood ratio statistic they highlight is most easily interpreted as pertaining to a hypothesis that is not generally of forensic interest. Specifically, it follows from the hypothesis that the observed sequencing reads were derived from biological material from an individual who is included as a member of the reference database used to run the method.

`IBDGem’s` likelihood ratios can also be interpreted as estimates of the standard forensic likelihood ratio. However, we show that with the numbers of markers and reference database sizes used by Nguyen and colleagues, the likelihood ratios produced by `IBDGem` are often very poor estimates, sometimes many orders of magnitude larger than ones that result from tests of the conventional defense hypothesis. If used in forensic work, these `IBDGem` likelihood ratios could potentially produce an impression of much stronger evidence for identity than is warranted. We suggest directions that might lead to likelihood ratios more in line with standard forensic use.

## Results

### The IBDGem approach

Typically, forensic likelihood ratios for identity take the form

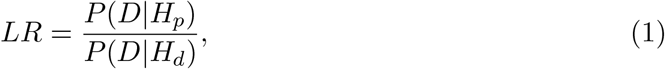

where *D* represents the event of observing the data (i.e. the evidence), *H*_*p*_ is a hypothesis advanced by the prosecution, and *H*_*d*_ is a competing, alternative explanation for the observed data friendly to the defense (Williams & Maskell, 2021). In identity testing scenarios, this general expression often appears as

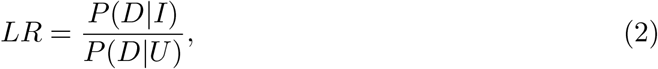

where *I* represents the event that the biological sample comes from the person of interest, and *U* represents the event that the biological sample comes from a person drawn at random from the population of alternative sources, unrelated to the person of interest. (For brevity, we refer to the population of alternative sources as “the relevant population” or just “the population” below.)

`IBDGem` aims to arrive at a likelihood ratio of the form in equation 2, taking as input sequencing reads, and in particular, the alleles that map to sites that are known to be biallelic. At the *i*th biallelic site, it is assumed that a person of interest may have one of three genotypes: (0,0), (0,1), or (1,1), where “0” represents a reference allele and “1” an alternate allele. A biological sample may produce a set of reads that map to this site, and the data from those reads can be represented as a pair of integers (*D*_*i*0_, *D*_*i*1_), where *D*_*ij*_ represents the number of alleles of type *j* observed at biallelic site *i*. (`IBDGem` assumes that all reads are of one of the two alleles at the biallelic locus.) All of our comments concern likelihood ratios for a single “analysis window”—a set of sites analyzed jointly by `IBDGem`. An analysis window is assumed to cover one contiguous genomic region containing a user-specified number of variable sites from which reads have been obtained. Nguyen and colleagues often use analysis windows of 200 variable sites.

`IBDGem` needs to compute two probabilities to form a likelihood ratio within a single user-defined window of variable loci: the probability of obtaining the observed sequencing reads given that the source of the reads (i) is the person of interest, and (ii) is an unrelated person from the population. If the source is the person of interest, whose genotype is assumed known with certainty, then the probability of observing a set of reads can be understood as arising from randomness in which of the person’s two allelic copies appear in the reads, and randomness from sequencing errors. If sequencing errors are symmetric and occur with probability *ϵ* per base pair, if each allele is equally likely to appear in a sequencing read, and if randomness in the total number of sequencing reads at a site is ignored, then (Nguyen et al., 2023),

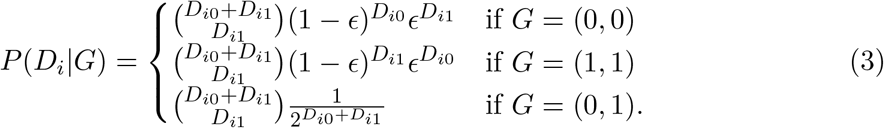

Nguyen and colleagues’ equations omit the binomial coefficient from the cases in which *G* = (0, 0) or *G* = (1, 1). However, this seems to be a typographical error, and the binomial coefficient is appropriately included in the `IBDGem` code.

`IBDGem` uses a default value of 0.02 for the (symmetric) sequencing error rate *ϵ*, though the user can choose other values. If the reads are independent (e.g. because variable sites are distanced enough to appear on different sequencing reads), then the probabilities produced by equation 3 for each of *k* variable sites within a single analysis window can be multiplied to form a probability of observing the sequencing data at the set of sites:

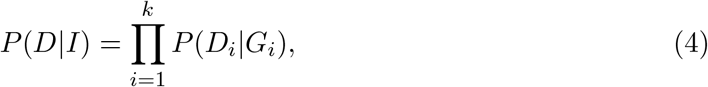

where *G*_*i*_ represents the genotype of the person of interest at site *i*, and *I* is the event that the biological sample comes from the individual of interest. Equation 3 implies that important contributors to this likelihood are the number of reads inconsistent with the person-of-interest genotype and the sequencing error rate. The likelihood is very small if there are many reads inconsistent with the person-of-interest genotype and the sequencing error rate is not large.

If a variable site from which reads have been obtained can be assumed to be in Hardy– Weinberg equilibrium with known allele frequency *p* in the relevant population, then one can compute the probability that a randomly chosen, unrelated individual would produce the observed reads at site *i* by summing *P* (*D*_*i*_|*G*) over the three possible genotypes, weighted by their population frequencies. That is, one can write

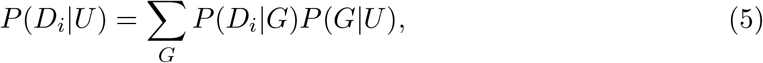

where the probabilities of the genotypes *P* (*G*|*U*) are the Hardy–Weinberg frequencies *p*^2^, 2*p*(1 *− p*), and (1 *− p*)^2^. If, further, genotypes at the variable sites are all statistically independent (that is, they are in linkage equilibrium), and we retain the assumption of independence of reads as before, then the likelihood of the data within a single analysis window can be computed by the product rule, giving

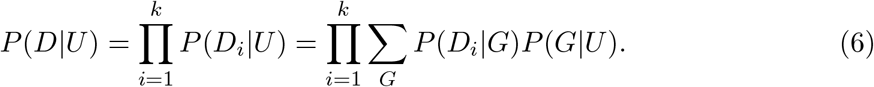

However, equation 6 will not give the correct probability if variable sites in linkage disequilibrium (i.e. sites with non-independent genotypes) are used. In particular, equation 6, by treating each site as statistically independent, risks overstating the amount of identifying information available at each site. Rather than providing a likelihood, it provides a composite likelihood (Larribe & Fearnhead, 2011). In dealing with closely spaced markers derived from sequencing, linkage disequilibrium is ubiquitous if sites are not pruned.

In response to the significant challenge presented by linkage disequilibrium, the makers of `IBDGem` developed and recommend an “LD mode” as a method to account for linkage disequilibrium. The LD mode approach is featured in Figure 1 of Nguyen and colleagues (2023) as a major advance of the method. Nguyen and colleagues suggest that the likelihood ratios arising from LD mode can be treated as testing the null hypothesis we label *U*, and they label “*IBD*0,” writing “`IBDGem` evaluates the likelihood of observing the sequence data if a test individual, whose genotype is known, was the source versus the likelihood of observing that same data if an unrelated individual was the source.” However, as we show below, the `IBDGem` approach to LD correction is most easily interpreted in general as testing a different null hypothesis, sometimes leading to statements that are more anticonservative even than those that follow from ignoring linkage disequilibrium.

**Figure 1:**
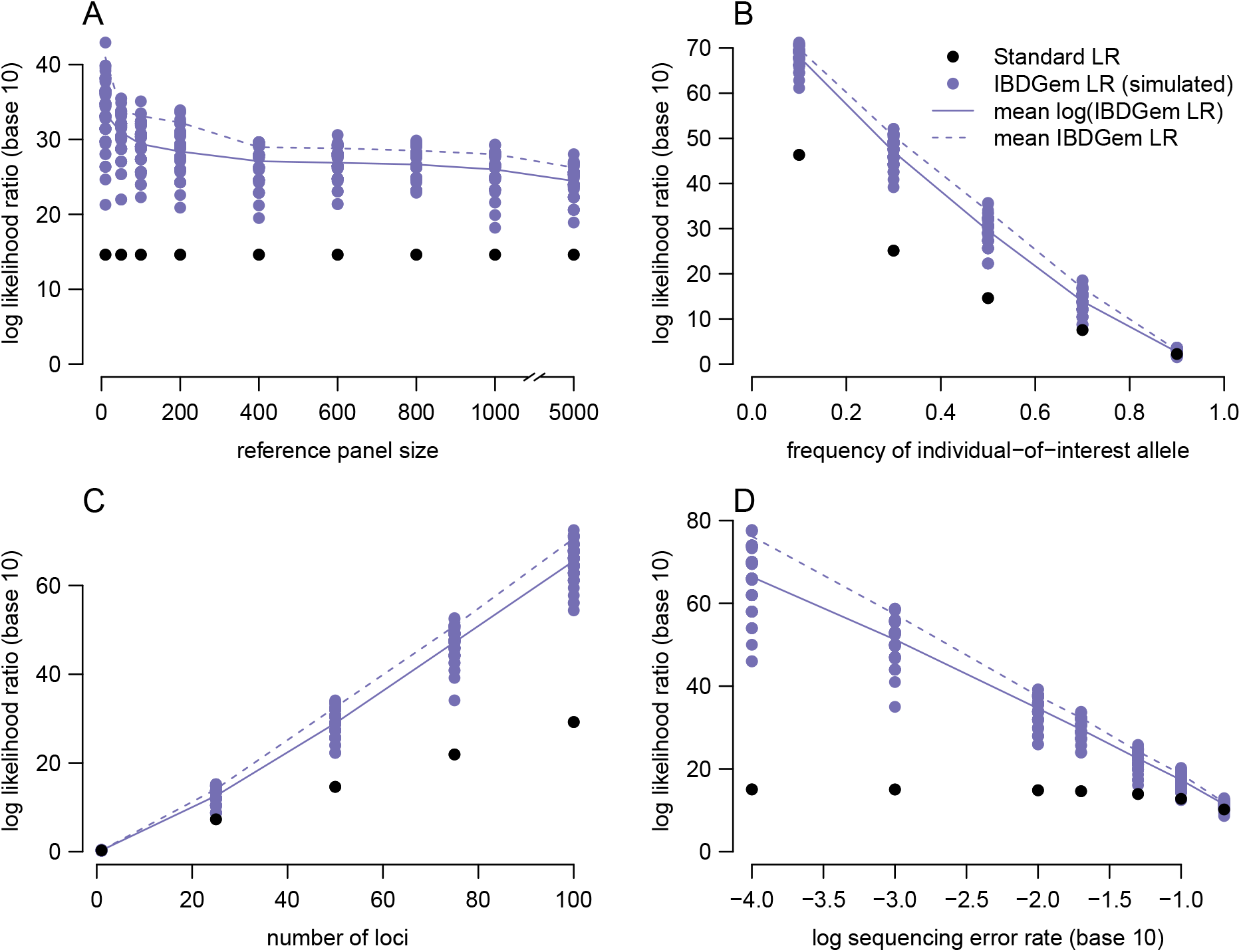
Dependence of the standard likelihood ratio (*P* (*D*|*U*) in the denominator) and `IBDGem` likelihood ratio (*P* (*D*|*R*) in the denominator) on parameters in the haploid model. Baseline parameters are reference database size *n* = 100, number of loci *k* = 50, allele frequency *p* = 1*/*2, and sequencing error rate *ϵ* = 0.02. One parameter is varied per panel: (A) reference database size, (B) allele frequency (axis label is 1 *− p*), (C) number of loci, (D) sequencing error rate. All likelihood ratios are displayed on a log scale (base 10). Standard forensic likelihood ratios are displayed in black, those from the `IBDGem` LD-mode approach in purple. There are multiple `IBDGem` dots per parameter value because LD-mode likelihood ratios depend on the specific individuals included in the reference database. For `IBDGem` likelihood ratios, dots represent values from simulated reference databases (100 per parameter configuration), solid lines give the mean of the log likelihood ratios, and dashed lines give the log of the mean likelihood ratios.

`IBDGem`’s LD mode is based on comparing reads with the specific individuals included in its reference database, rather than comparing them with allele or haplotype frequencies estimated from the reference database. The stated rationale is that in comparing with the specific individuals in the reference database, `IBDGem` “asks how the sequence data look against random individuals who we know share zero IBD chromosomes” (Nguyen et al., 2023). If one has access to a reference database of *n* individuals with known genotypes, then one can compute the probability in equation 4 with respect to each of these *n* individuals, simply replacing the person of interest’s genotypes with the genotypes of each member of the reference database in turn. Nguyen and colleagues (2023) then compute the average of this probability over individuals in the reference database,

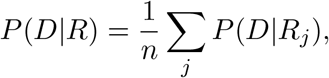

where *R*_*j*_ represents the hypothesis that the *j*th member of the reference database is the source of the reads. Nguyen and colleagues refer to this quantity as an estimate of *P* (*D*|*IBD*0) that accounts for LD, and they use it as the denominator in reported likelihood ratios. (They use *IBD*0 to denote the event that the source is an individual with no identity by descent segments with the person of interest in the region being examined.) We, however, label this quantity *P* (*D*|*R*), the probability of the data arising from a (uniformly) randomly chosen member of the reference database. (*R* denotes the event that the source is an individual drawn uniformly at random from the reference database.) This is because it has exactly that interpretation. By the law of total probability, the probability of observing the given reads from a random member of the reference database is the sum of the products of the probability of selecting each specific member of the reference database multiplied by the probability of observing the reads from that member of the reference database. Expressing the law of total probability in this case leads directly to the probability computed by `IBDGem`:

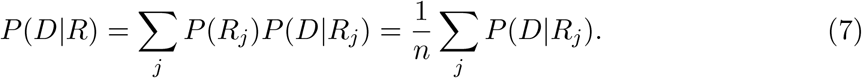

Thus, the value of *P* (*D*|*IBD*0) computed by `IBDGem`’s LD mode—that is, the average, across the multilocus genotypes in the reference database, of the probability of observing the reads—is equal to the probability (under their model for sequencing reads) of observing the data from a randomly selected member of the reference database. This probability is not generally equal to the one of forensic interest—that of observing the data from a random, unrelated individual in the population of alternative sources—unless either (i) the individuals included in the reference database are exactly the same individuals in the population of alternative sources or (ii) the multilocus genotype frequencies in the reference database are the same as in the relevant population of alternative sources. (Note that for (i), the condition is that the reference database and the population of alternative sources are identical, not just that the reference database is composed of members of the population of alternative sources. And for (ii), it is the *multilocus* genotype frequencies that matter, not the marginal frequencies of the genotypes in the window—more on this below.) Nguyen and colleagues do not point out the difference.

Thus, whereas the likelihood ratio generally of forensic interest is, in our notation, *P* (*D*|*I*)*/P* (*D*|*U*), `IBDGem`’s LD mode forms a likelihood ratio

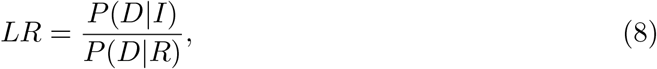

replacing the probability of observing the data from an unrelated person in the relevant population, *U*, with the probability of observing the data from a random person in the reference database, *R*. One way to express the difference between these likelihood ratios is that in typical forensic practice, chance variation due to both genotyping error and genetic variation is considered. Existing likelihood ratios in forensics posit a model for combinations of genotypes that are not observed in the reference database (Balding & Steele, 2015; Buckleton et al., 2016; Butler, 2010). However, in the likelihood ratio produced by `IBDGem`’s LD mode, there is no model for genetic variation outside the reference database, at least as concerns sites within the same analysis window.

To generalize from the reference database to the population of potential sources of the sample requires a model of genetic variation, not just a model of sequencing reads and errors therein. A very low value of *P* (*D*|*R*) can be informative about identity, particularly if the reference database is large, but it cannot generally be interpreted as the probability of observing the data from a randomly chosen, unrelated person from the relevant population, *P* (*D*|*U*).

One way in which *P* (*D*|*U*) and *P* (*D*|*R*) differ can be seen immediately: if the genotyping error rate is set to 0, and read data have at least one inconsistency with every member of the reference database, then *P* (*D*|*R*) = 0. This is true even for arbitrarily small reference databases and arbitrarily few loci, even though a small reference database with few loci represented leaves open the possibility that many individuals in the population carry genotypes consistent with the observed reads, meaning that *P* (*D*|*U*) *>* 0. Although this hypothetical analysis is not realistic—the authors of `IBDGem` do not recommend setting the error rate to zero—it shows the difference between standard forensic likelihood methods and `IBDGem`. In `IBDGem`, the reads within an analysis window that are incompatible with the full (i.e. multilocus) genotype of each specific individual in the reference database can only be explained by sequencing error. Another way to understand this hypothetical is that `IBDGem`’s LD mode relies on the assumption that the average likelihood among reference individuals is a good proxy for the likelihood of the data in the population at large. However, if the assumed sequencing error is low, most individuals in the population will have very low (or, in this hypothetical case, zero) likelihoods. However, there may in fact be a fraction of individuals in the population whose genotypes explain the data well and thus have high likelihoods. If none of these latter individuals are included in the reference, then *P* (*D*|*R*) will be very small, potentially smaller than the population value by many orders of magnitude, leading to overstated likelihood ratios for identity.

We illustrate that the values of *P* (*D*|*U*) and *P* (*D*|*R*)—that is, the probability of observing the data from an unrelated member of the population of interest and the average probability of observing the reads from the reference individuals—can diverge widely within an analysis window in examples below, often leading to likelihood ratios that differ by many orders of magnitude.

### A simplified haploid example

For illustration, we focus on a simplified case. A haploid individual of interest carries “0” alleles at each of *k* biallelic sites. At each of these *k* sites, the frequency of the “1” allele is *p*, and alleles at distinct sites are independent. We have a reference database of *n* randomly selected individuals with known alleles at each of the *k* sites. We also have sequencing reads from a biological source, exactly one read from each site, with assumed symmetric error rate *ϵ*, and all of the reads are “0” alleles, meaning that they are consistent with the individual of interest.

In this scenario,

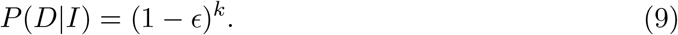

That is, if the haploid of interest is the source of the sequencing reads, we will observe sequencing reads that match the haploid of interest at all *k* sites if and only if there are no sequencing errors at any of the *k* sites.

We can also compute the probability of observing the reads from an individual with *L* = *l* “1” alleles and *k−l* “0” alleles as *P* (*D*|*L* = *l*) = *ϵ*^*l*^(1 *−ϵ*)^*k−l*^. That is, if an individual does not match the sequencing reads at *l* positions, then to produce the sequencing reads, there must have been exactly *l* sequencing errors, occurring in exactly the positions of mismatch.

The probability of observing the sequencing reads from an unrelated individual in the population can be obtained by summing *P* (*D*|*L* = *l*) over the distribution of *L* for unrelated individuals,

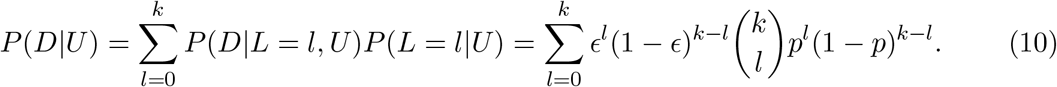

The second step follows because under the described model, *L* has a Binomial(*k, p*) distribution among individuals unrelated to the haploid of interest, giving 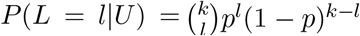. The sum in equation 10 is clearly bounded from below by its *l* = 0 term, (1™*p*)^*k*^*(1™ϵ)*^*k*^, the probability that an individual unrelated to the haploid of interest has a genotype exactly matching the haploid of interest and there are no sequencing errors. Thus, the likelihood ratio formed by dividing equation 9 by equation 10 is bounded from above by 1*/*(1 *− p*)^*k*^. An alternative, equivalent expression for *P* (*D*|*U*) closer to the spirit of equation 6 is

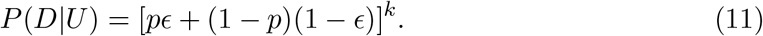

In contrast, the probability of observing the data from a randomly chosen member of the reference database is sensitive to the genotypes of the specific individuals in the reference database. For illustration, Table 1 shows a randomly drawn reference database with *n* = 5 individuals, *k* = 10 loci, and *p* = 1*/*2 the allele frequency at each locus. These hypothetical individuals were simulated using exactly the assumed allele frequencies. For these parameters and a sequencing error rate of *ϵ* = 0.02, *P* (*D*|*I*) *≈* 0.82, and *P* (*D*|*U*) *≈* 9.8 × 10^−4^. The typical forensic likelihood ratio comparing the probability of the observed data under the hypotheses that the sample source is the person of interest vs. a random individual from the population is thus *P* (*D*|*I*)*/P* (*D*|*U*) *≈* 837. However, the probability of observing the data from a randomly chosen individual in the reference database in Table 1 is only *P* (*D*|*R*) *≈* 1.2 × 10^−9^. Thus, in this example, using *P* (*D*|*R*) as a stand-in for *P* (*D*|*U*), as `IBDGem` does in LD mode, gives a likelihood ratio of *≈* 6.9 × 10^8^, a difference of over five orders of magnitude.

**Table 1:** Simulated data for the haploid model with *n* = 5 and *k* = 10. Individuals are in rows, loci in columns.

|  |  |  |  |  |  |  |  |  |  |
|---|---|---|---|---|---|---|---|---|---|
| 0 | 1 | 1 | 1 | 0 | 1 | 1 | 0 | 1 | 0 |
| 0 | 1 | 1 | 1 | 1 | 0 | 1 | 0 | 1 | 0 |
| 1 | 1 | 1 | 0 | 1 | 0 | 0 | 0 | 1 | 0 |
| 1 | 1 | 0 | 1 | 0 | 0 | 1 | 0 | 1 | 0 |
| 0 | 0 | 1 | 1 | 1 | 1 | 0 | 1 | 0 | 1 |

In this hypothetical example, there is a large difference between the likelihood ratios formed by the quantities we label *P* (*D*|*U*), used in standard forensic practice, and *P* (*D*|*R*), computed by `IBDGem`’s LD mode. We show by simulation that similarly large or larger differences can occur in other scenarios. Figure 1 shows how the two likelihood ratios and their relationship change with the number of loci, size of the simulated reference database, and error rate. For many sets of parameter values, likelihood ratios of the form `IBDGem` uses are many orders of magnitude larger than standard forensic likelihood ratios.

The difference between the likelihood ratios is determined by *P* (*D*|*U*) and *P* (*D*|*R*)—if the probability of obtaining the observed reads from a random member of the reference database is much lower than the probability of obtaining the observed reads from a random member of the relevant population, then the likelihood ratio produced by `IBDGem`’s LD mode will be much larger than the likelihood ratio testing the hypothesis generally of forensic interest. If no one in the reference database closely matches the reads produced, then the only way to accommodate the data with the reference database is to posit sequencing errors. Thus, the `IBDGem` likelihood ratios will be especially large when the assumed sequencing error rate is low (or, for the same set of variable sites, when sequencing depth is high), when the observed reads contain alleles from rare haplotypes (less likely to be observed in the reference database), or when the reference database is small (meaning that it is unlikely to contain haplotypes similar to the observed reads). We see that higher sequencing error rates (in combination with relatively low sequencing depth) cause the two likelihood ratios to converge at lower reference sample sizes, because mismatching alleles can be readily explained as sequencing errors.

### IBDGem LD-mode likelihood ratios with simulated diploid data

We next extend the haploid model used for illustration to diploids and replicate the basic patterns seen in our haploid implementation using `IBDGem` itself. We simulate diploids by pairing together two haplotypes of sites in linkage equilibrium with allele frequencies obeying the expected site frequency spectrum for a neutrally evolving population of constant size. In these simulations, the individual of interest no longer carries “0” alleles at each of the *k* biallelic sites—they now carry a mix of heterozygous and homozygous genotypes. Under this model, we simulate sequencing reads for the target individual on the basis of their genotype while including sequencing errors. We use the same error rate used to simulate sequencing reads to analyze the data in `IBDGem`, using the –error-rate flag. We also simulate a reference database of *n* diploid individuals with known genotypes at each of the *k* loci. See Methods for more description.

We use the simulated sequencing reads, the genotypes of the target individual and reference database, and allele frequencies as input to `IBDGem`, running it in both LD and non-LD modes. Because we are simulating genotypes in linkage equilibrium, the non-LD-mode likelihood ratio is correct and equal to the standard forensic likelihood ratio. Figure 2 shows how within a single analysis window, `IBDGem`’s LD-mode and non-LD-mode likelihood ratios and their relationship change with the size of the simulated reference database, the sequencing depth (i.e. number of reads per variable site), the number of loci, and error rate. The basic patterns are the same as in the haploid case—for many sets of parameters, the likelihood ratios produced by `IBDGem`’s LD mode are much larger than the likelihood ratios produced by the non-LD mode. Similar patterns are seen with more realistic allele-frequency distributions including the CEU and YRI subsets of the 1000 Genomes panels (Auton et al., 2015) (Supplementary Fig. S1-S2).

**Figure 2:**
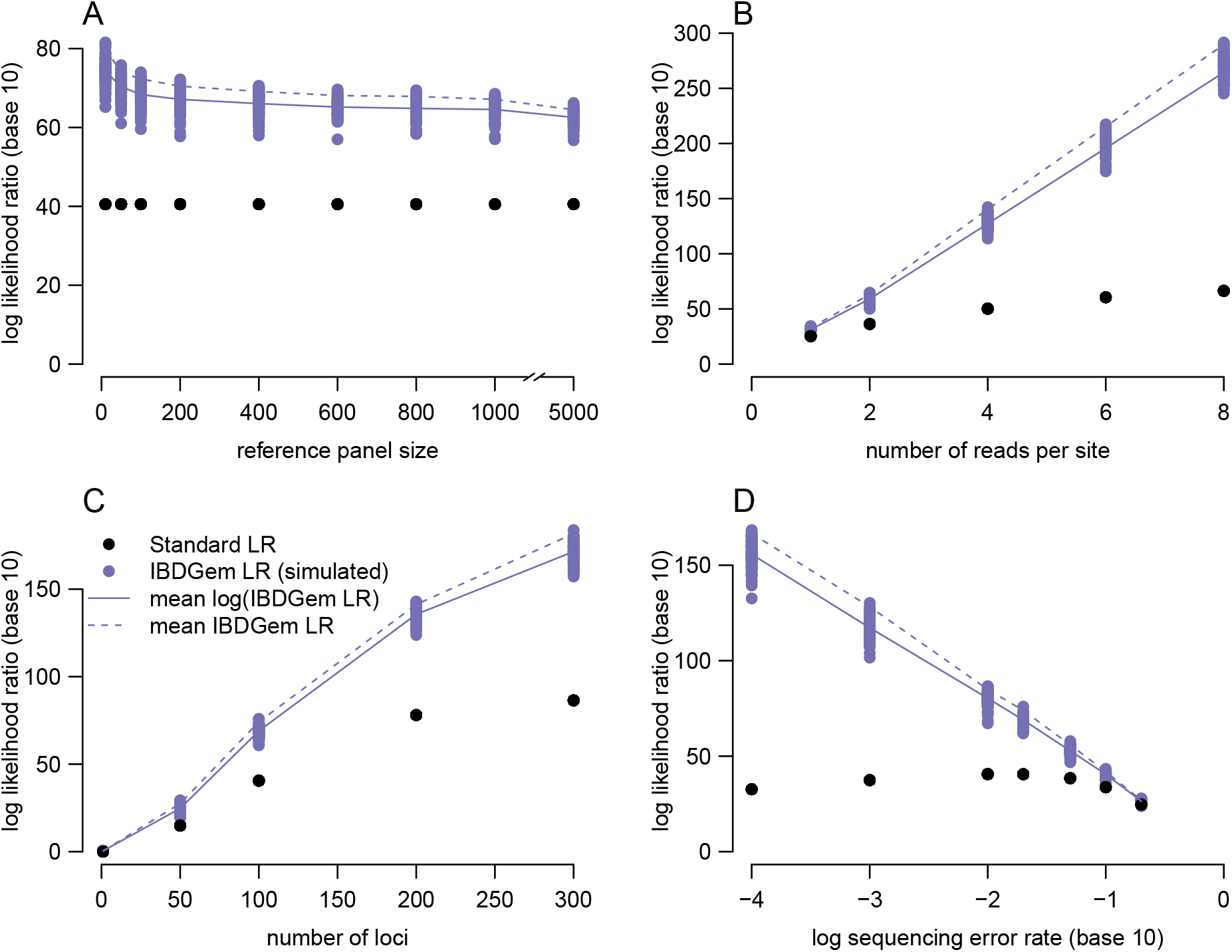
Dependence of the standard likelihood ratio (*P* (*D*|*U*) in the denominator) and `IBDGem` likelihood ratio (*P* (*D*|*R*) in the denominator) on parameters in a diploid setting obeying the neutral site frequency spectrum with no linkage disequilibrium. These likelihood ratios were computed using `IBDGem` itself. Baseline parameters are reference database size *n* = 100, number of loci *k* = 100, sequencing error rate *ϵ* = 0.02, and exactly two reads per site. One parameter is varied per panel: (A) reference database size, (B) coverage depth, (C) number of loci, (D) sequencing error rate. All likelihood ratios are displayed on a log scale (base 10). Likelihood ratios computed using `IBDGem` non-LD mode—which are correct in this case, as the simulated genotypes are not in LD—are displayed in black, those from the `IBDGem` LD mode approach in purple. For LD-mode likelihood ratios, dots represent values from simulated reference databases (100 per parameter configuration), solid lines give the mean of the log likelihood ratios, and dashed lines give the log of the mean likelihood ratios.

### IBDGem LD-mode likelihood ratios under linkage disequilibrium in haploids

`IBDGem`’s LD mode was developed in order to account for correlations among loci, or linkage disequilibrium (Slatkin, 2008). Linkage disequilibrium is ubiquitous in sequence data, which may produce reads from closely spaced loci.

We ran simulations to explore the effect of linkage disequilibrum on likelihood ratios computed in the manner of `IBDGem`, extending the haploid model above. We use two different simplified representations of linkage disequilibrium, one that contains blocks of markers in perfect (*r*^2^ = 1) linkage disequilibrium, and one in which all markers are in complete (*D*^*′*^ = 1) linkage disequilibrium, but no markers are in perfect (*r*^2^ = 1) linkage disequilibrium. The models of linkage disequilibrium we consider are not chosen to be realistic descriptions of LD in humans. Instead, we have chosen them because they allow easy computation of the standard forensic likelihood ratio, which we can then compare with the numerical values produced by `IBDGem`’s LD mode.

Specifically, to simulate blocks of perfect linkage disequilibrium, we imagine that every variable locus has *m* exact copies in the same “block”—that is, if a “1” allele is observed in a block, then all other alleles will be “1” alleles, and similarly for “0” alleles. In this scenario, the no-LD model will count distinct markers in the same block as independent pieces of information, when the genotypes are in fact completely redundant. `IBDGem`’s LD mode was developed to account for linkage disequilibrium in scenarios like this one, avoiding the redundancy in the default mode’s analysis of markers that may be in linkage disequilibrium.

We also simulate a scenario in which *k* markers are in complete, but not perfect, linkage disequilibrium, forming *k* + 1 distinct possible haplotypes at equal frequency. The specific scheme we use is consistent with a “caterpillar” gene tree (Steel, 2016), and is exemplified in Table 2 for the case of *k* = 4 variable loci. In this scheme, all loci are in complete linkage disequilibrium (*D*^*′*^ = 1), meaning that variation in the population is compatible with a history in which no recombination has occurred, but because the allele frequencies differ, they are not in perfect linkage disequilibrium (VanLiere & Rosenberg, 2008). For more information, see Methods.

**Table 2:** Five possible haplotypes in the haploid “caterpillar” simulation with *k* = 5 loci, along with the haplotype frequencies (1st column) and implied allele frequencies (bottom row).

|  |  |  |  |  |
| --- | --- | --- | --- | --- |
| 1/5 | 0 | 0 | 0 | 0 |
| 1/5 | 1 | 0 | 0 | 0 |
| 1/5 | 1 | 1 | 0 | 0 |
| 1/5 | 1 | 1 | 1 | 0 |
| 1/5 | 1 | 1 | 1 | 1 |
| $p$ | 4/5 | 3/5 | 2/5 | 1/5 |

Figure 3 shows results from these simulations. As expected, `IBDGem`’s non-LD approach produces biased likelihood ratios in the presence of linkage disequilibrium. But, as in the other scenarios, LD mode does not generally correct this problem if the reference database size is not large enough that it would be expected to contain the exact haplotype of the individual of interest.

**Figure 3:**
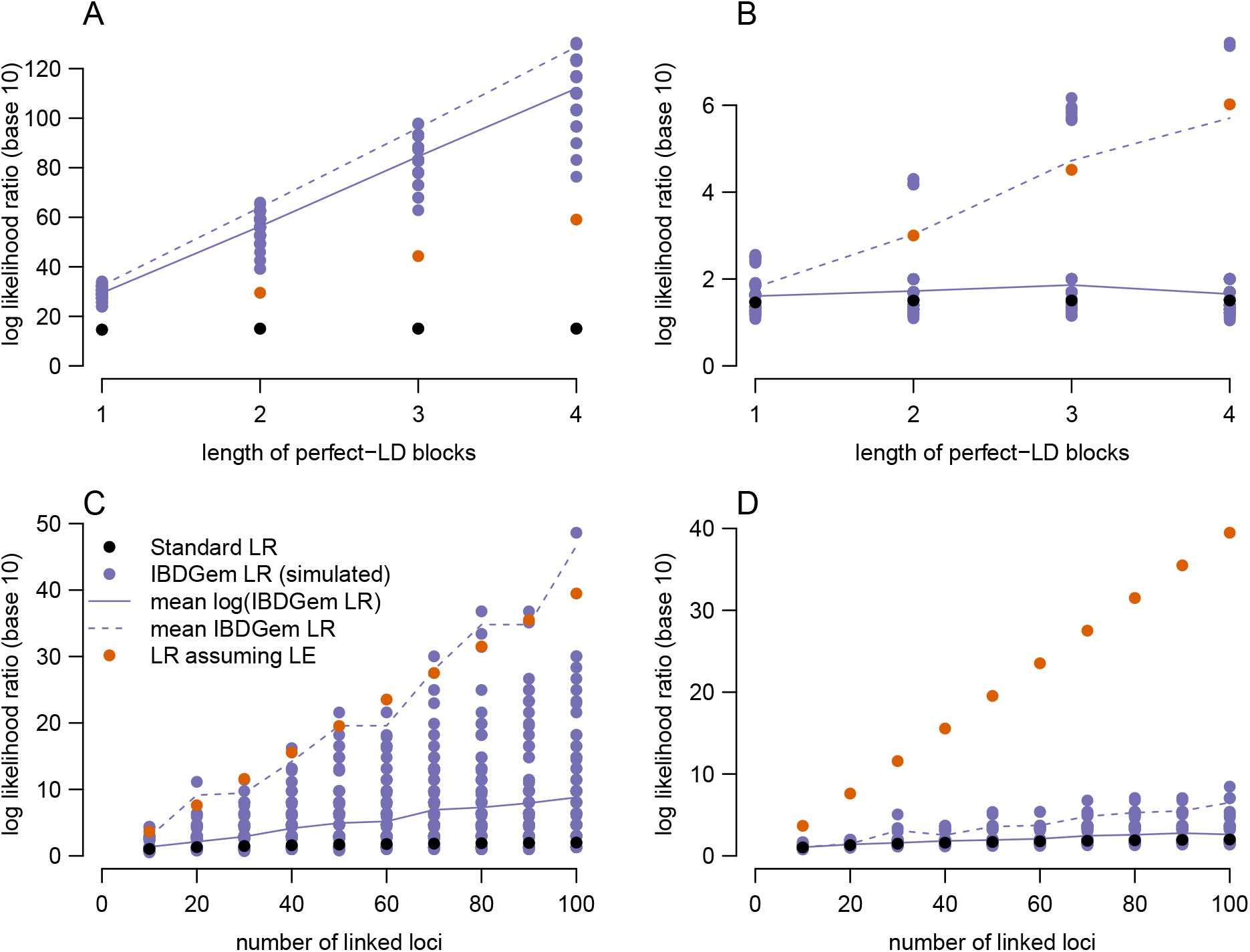
In the presence of linkage disequilibrium, likelihood ratios following the model for `IBDGem`’s non-LD mode will overstate evidence for identity. However, as in the other examples, LD mode only gives roughly correct likelihood ratios reliably when the reference database size is considerably larger than one over the frequency of the haplotype of interest. Conventions are the same as in Figure 1, but the orange dots indicate likelihood ratios computed using the method of `IBDGem`’s non-LD mode, which are now incorrect due to the LD in the simulated genotypes. Panels A and B show results from perfect blocks of LD, as described in the text, and panels C and D show results from the caterpillar haplotype model exemplified in Table 2. In all panels, the sequencing error rate is *ϵ* = .02, and in panels A and B, *p* = 0.5 and *n* = 100. In Panel A, *k* = 50 and is now interpreted as the number of sets of mutually independent loci, whereas in Panel B, *k* = 5. In Panel C, *n* = 20, whereas in Panel D, *n* = 100.

In real human data, linkage disequilibrium is likely to be intermediate between the cases of linkage equilibrium and of complete or perfect LD that we simulated. We chose to simulate under these simplified models because they render the standard forensic likelihood easily tractable. In neither case does `IBDGem` LD mode recover the standard forensic likelihood ratio.

### IBDGem LD-mode likelihood ratios under linkage disequilibrium in diploids

We extended the above model exploring perfect and complete linkage disequilibrium to diploids. More details on the implementation are in the Methods section. We observed similar patterns to the haploid case – likelihood ratios generated under both `IBDGem`’s non-LD and LD modes produce biased likelihood ratios in the presence of linkage disequilibrium (Supplementary Fig. S3). Increasing the reference database size does not necessarily correct the problem unless the analysis window spans a small number of variable sites (Supplementary Fig S4).

*P* (*D*|*R*) and *P* (*D*|*U*) are equivalent if the joint frequencies of all within-window genotypes in the panel are exactly as they are in the relevant population, meaning that the `IBDGem` likelihood ratios might be interpreted in the same way as standard forensic likelihood ratios if this condition is met, or perhaps as estimates of the standard likelihood ratio if the condition is approximately met. However, the condition is difficult to meet with reference database sizes substantially smaller than the relevant population itself, so long as there are at least a moderate number of markers in the analysis window that are not in complete LD. For example, if there are 30 variable diploid markers that are not completely linked in an analysis window, then there are 3^30^—that is, approximately 2 × 10^14^—possible multilocus genotypes, orders of magnitude more than the number of humans in the world. Most of these will necessarily not exist in the relevant population. However, there will also be many multilocus genotypes that exist in the relevant population at low frequency—indeed, when this many markers are considered with appreciable minor allele frequencies, nearly all multilocus genotypes are low frequency—but are not included in reference databases that are not approximately as large as the population, such as the 1000Genomes panel. Thus, many multilocus genotypes that have nonzero frequency in the population will have zero frequency in the reference database, leading to the large likelihood ratios seen in our simulations.

Supplementary figure S5 shows that for moderately sized windows, nearly every multilocus genotype in 1000Genomes is unique, meaning that samples in the thousands are not nearly large enough to include all multilocus genotypes, let alone to quantify their frequencies.

`IBDGem` is robust to diploid phase, meaning it implicitly covers the haplotypes that would be captured by a haplotype-based, phase-sensitive method that could be produced by combining the alleles (within an individual) into haplotypes in any way. However, this robustness to phase does not allow for combinations of alleles sampled from haplotypes that appear in different individuals.

This limitation means that even if there is a pair of haplotypes in the reference database that easily explains the sequencing reads, `IBDGem` will still produce a very small likelihood that the data come from a person unrelated to the person of interest if those haplotypes do not appear in the same individual. Figure 4 compares the log-10 likelihood ratios produced by `IBDGem`’s LD mode in two cases. Either there is an individual in the reference database carrying two haplotypes from which all the simulated sequencing reads come (unfilled circles), or the two haplotypes from which the reads come appear in different individuals (filled circles). When the haplotypes that explain the reads appear in the same reference individual, the likelihood ratios provide only weak evidence for identity. However, when these haplotypes appear in two different individuals, the likelihood ratios are over ten orders of magnitude larger and in a range that would typically be considered strong evidence of identity, despite the fact that there are haplotypes in the reference database that easily explain the reads. We also note that shuffling haplotypes among individuals does not change the LD at the level of haplotypes.

**Figure 4:**
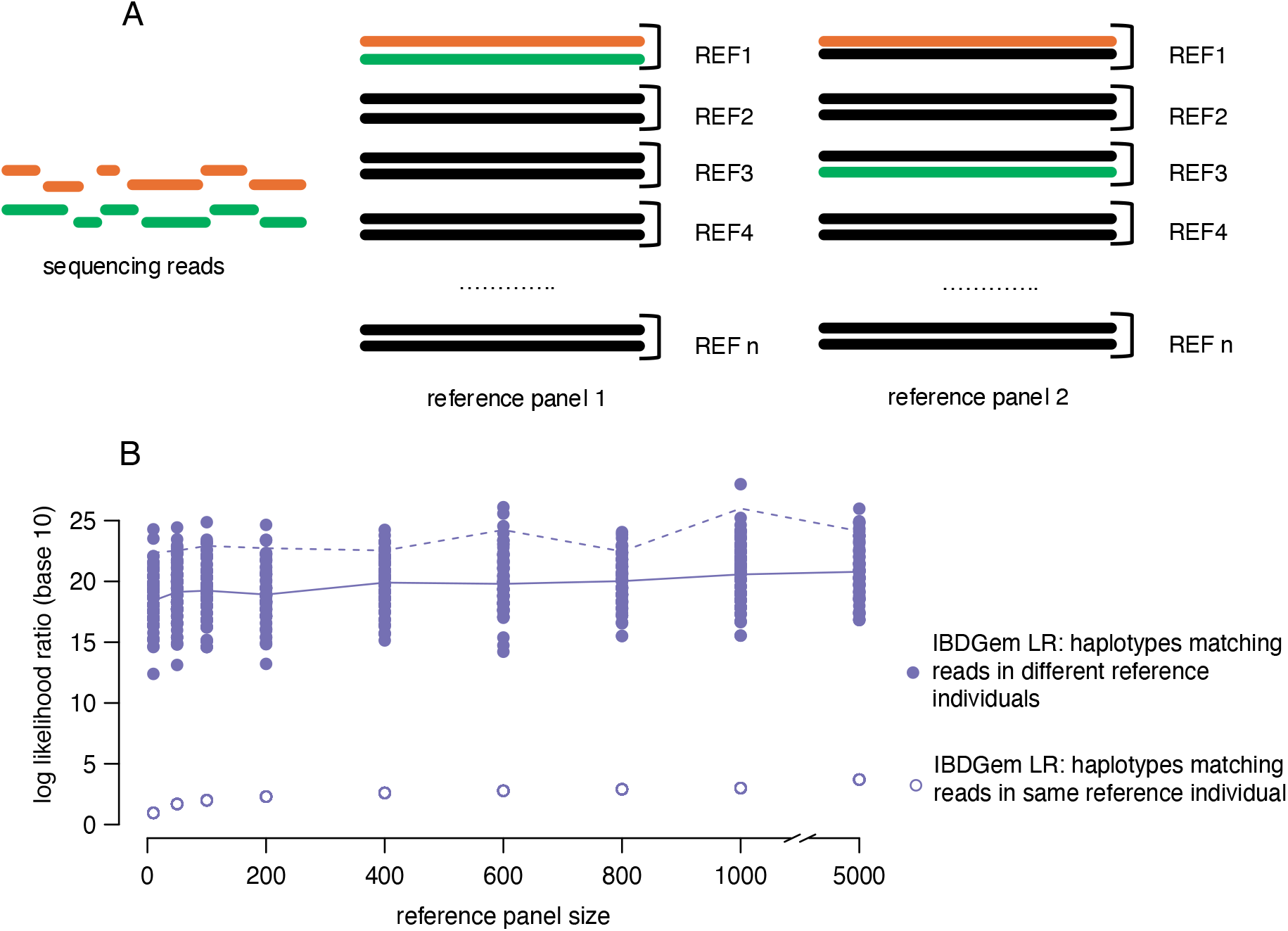
`IBDGem` is restricted to haplotypes from a single reference individual, excluding any consideration of haplotype combinations across individuals. Panel A displays a schematic of the experiment performed here: a set of sequencing reads is analyzed by `IBDGem` with two different reference databases containing the same haplotypes. However, in one analysis, the haplotypes from which the sequencing reads come appear in the same reference individual, and in the other analysis, they appear in different individuals. Panel B displays the base-ten log-likelihood ratios that result from the two analysis performed on 100 different simulated reference databases of each of several sizes. Parameters are as in Figure 2: the sequencing error rate is *ϵ* = .02, the number of loci *k* = 100, and there are two reads per site.

We carried out further simulations in which we rearranged the haplotypes within the reference database to form new individuals, while preserving the same haplotypes but not forcing the appearance of haplotypes that explain the reads. In doing so, the likelihood ratio often changes by several orders of magnitude (2-3 orders of magnitude at median and over ten orders of magnitude in the most extreme cases), since `IBDGem`’s LD-mode does not consider combinations of haplotypes across individuals (Supplementary Fig. S6).

### Recommended practices for using IBDGem

If the reference database contains the entire population that is the target of inference— or alternatively, if the multilocus genotype frequencies in the reference database exactly match those in the population of interest—then *P* (*D*|*R*) and *P* (*D*|*U*) become equivalent, and `IBDGem`’s LD mode likelihood ratio addresses the forensic question of interest. We might expect, similarly, that if the reference database is large enough to contain individuals with multilocus genotypes consistent with those of the person of interest at their frequency in the relevant population, then *P* (*D*|*R*) and *P* (*D*|*U*) should be similar. In simulations with small numbers of loci, it is tractable to simulate reference databases with *n >>* 1*/P* (*D*|*U*), meaning that the individual of interest’s multilocus genotype is likely to be observed in the reference database. By using a small analysis window size, we reduce the number of unique within-window multilocus genotypes, which, in turn, allows the joint frequencies of within-window genotypes in the reference database to closely match those in the relevant population.

Figure 5 shows simulations similar to Figure 2, but with smaller numbers of loci. In panel A, likelihood ratios using *P* (*D*|*U*) and *P* (*D*|*R*) appear to match if the reference database is large enough that we would expect several copies of the individual-of-interest genotype in the reference database. In panel B, *P* (*D*|*U*) and *P* (*D*|*R*) appear to match if the number of loci considered within a single window of analysis is limited to a small number.

**Figure 5:**
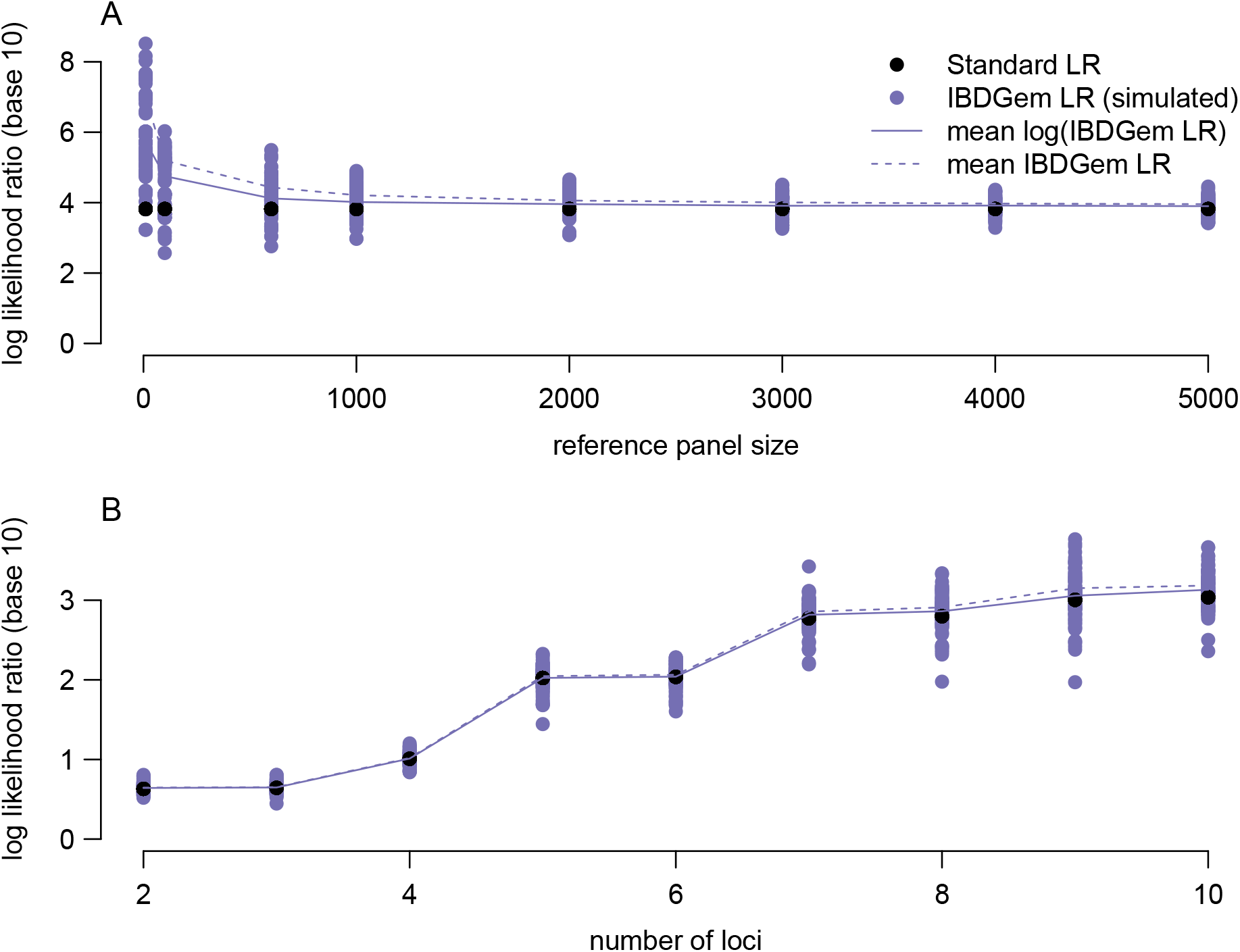
When the reference sample is large enough to include the individual-of-interest genotype, standard and `IBDGem` likelihood ratios roughly match. Conventions and most parameters are the same as in Figure 2, but here, the values of *k* are smaller. In panel A, the number of loci is *k* = 5, and in panel B, the reference database size is *n* = 100. In both panels, the sequencing error probability is *ϵ* = .02, there are two reads per site, sites are unlinked, and allele frequencies obey a standard neutral SFS.

## Discussion

We showed that the likelihood ratios produced by `IBDGem` within a single analysis window of variable loci can be badly misleading about the strength of evidence in identity inference in scenarios similar to those we have explored by simulation. Likelihood ratios produced by `IBDGem`’s non-LD mode can be strongly anti-conservative in the presence of linkage disequilibrium, as expected. As such, the makers of `IBDGem` developed LD mode “to account for linkage disequilibrium” by using a reference database and “compar[ing] the observed data against the genotypes of these unrelated individuals” (Nguyen et al., 2023). But this approach changes the meaning of the likelihood ratio, which Nguyen and colleagues do not note—the LD-mode likelihood ratios can be derived from the null hypothesis that the data come from a random member of a specific reference database, rather than a random, unrelated person from the population of potential sources of the sample.

In a forensic context, the defense’s hypothesis for explaining sequencing reads from a biological sample is generally that the biological sample comes from an unknown person in the population of potential contributors of the sample, not from a member of a specific reference database. Whether a member of the 1000 Genomes panel (Auton et al., 2015), for example, is the source of the biological sample is not generally the question of interest. Moreover, the likelihood ratios obtained from the LD-mode approach of `IBDGem` can differ from the ones calculated with respect to the standard forensic hypotheses by many orders of magnitude, potentially producing an impression of much stronger support for inclusion of a person of interest as the source of biological material than is warranted.

In our use of `IBDGem`, we noted that large reference database sizes tend to reduce the discrepancy between *P* (*D*|*U*) and *P* (*D*|*R*), and large numbers of markers tend to increase it. The reasons for these trends are related. `IBDGem`’s LD-mode likelihood ratios reproduce standard likelihood ratios if the reference-panel frequencies of all multilocus genotypes within an analysis window match the relevant population frequencies. We emphasize that it is the *multilocus* genotype frequencies that must match, not the marginal frequencies at each site. As the reference database size grows, the reference database frequencies approach the overall population frequencies, gradually bringing *P* (*D*|*U*) and *P* (*D*|*R*) closer together. However, if there are many markers that are not in complete LD within an analysis window, then there will be many different multilocus genotypes present in the population, making it much more difficult to estimate their frequencies. In particular, some joint genotype frequencies that are nonzero in the population may be zero in the reference database, potentially leading to values of *P* (*D*|*R*) much smaller than *P* (*D*|*U*) and thus large likelihood ratios.

In practice, `IBDGem` can be applied multiple times per genome, once for each of several user-defined windows of variable loci. Our results are with regard to a single analysis window. There are further questions about how to combine results across windows. It is worth mentioning that combination of results across windows could allow the consideration of some genetic variation that is not included in the reference database. For example, if a set of reads in one window matches reference individual 1, and in another window, matches reference individual 2, then the combination of the likelihood ratios from these two windows could indicate that the reads are easily explained by observed genetic variation, even if no single individual in the reference database matches the reads well in both windows. However, crucially, this is only true across windows, not within them. Some methods of combining information across windows may lead to conservative likelihood ratios in practice, even if the window-wise likelihood ratios are anti-conservative. To us, it seems preferable to begin from likelihood ratios that match the forensic hypothesis, rather than to devise a way of combining information across windows to undo any potential problems with the within-window likelihoods.

The work of Nguyen and colleagues (2023) establishes that there is substantial information relevant for identity inference even in low-coverage sequencing reads. The issue is attaching a reliable (or conservative) assessment of the strength of the evidence for identity. There are several possible routes by which approaches like that of `IBDGem` might pursue this goal. Perhaps the simplest—and the one taken by recent approaches to the same problem (Andersen et al., 2025; Mostad et al., 2023)—is to prune variable sites such that only sites in linkage equilibrium are likely to be included, and then use the product rule. Though this approach throws away information, a great deal is still left over. (However, linkage disequilibrium arising from population structure would be a key consideration, and it is not clear whether sufficient genotype information is currently available to adequately assess linkage disequilibrium for what are likely to be relevant pools of alternative sources of biological material.) A related approach would be to use short analysis windows—for example, with reference databases approximately as large as the 1000Genomes panel, it might be possible to estimate multilocus genotype frequencies well for sets of, say, 2-10 variable sites. (The exact number of variable sites that can be included in a window will depend on the reference database size, allele frequencies, and the degree of LD within the window. However, the 200-site windows suggested by Nguyen and colleagues are likely too large to estimate multilocus genotype frequencies accurately—and, in particular, to identify all the multilocus genotypes with nonzero frequency in the population of alternative contributors—with attainable sample sizes.)

A second approach would be to borrow methods from the analysis of rare “lineage markers,” such as Y-STRs (Brenner, 2010; Buckleton et al., 2011)—when no multilocus genotypes in a size-*n* reference database of randomly selected individuals can readily explain the observed reads, then the probability of the underlying multilocus genotype might be assigned a value based on 1*/n*, perhaps with some adjustment (e.g. the use of the lower bound of a 95% confidence interval for the frequency of a multilocus genotype not observed among *n* random individuals). Such an approach would be concordant with our result that *P* (*D*|*U*) and *P* (*D*|*R*) tend to be similar if the database is large enough to include multiple instances of the individual of interest’s haplotype. Along these lines, one might imagine that analysts using a program like `IBDGem` could focus on, instead of the average likelihood of the data among members of the reference, the fraction of members of the reference who produced likelihoods larger than the person of interest, along with the number and appropriateness of the reference individuals considered.

A third approach might be to use the reference data to build a model of haplotype structure (Li & Stephens, 2003) and use the estimated haplotype frequencies as a basis for inference. This approach is suggested, but not pursued, by Mostad and colleagues (2023). A haplotype model would also likely be the easiest to extend to analysis of mixed samples.

The issues we point out here regarding the interpretation of the likelihood ratio arise even in panmictic populations. More work is needed to understand the effects of population structure and representation in the database (Zavala et al., in press), as well as hypotheses involving biological relatives of the person of interest. Structure will lead to departures from Hardy–Weinberg equilibrium and from linkage equilibrium, even for unlinked markers, causing model violations for `IBDGem`’s non-LD mode. Further, if the source of the biological material in an evidence sample is from a population not well represented in the reference database, then the evidence against a person of interest may be overstated, a recurrent issue in forensic genetics (Balding & Nichols, 1994; Lewontin & Hartl, 1991). Given the strong dependence of `IBDGem`’s LD mode on the specific individuals included in the reference database, issues of population structure and representation may be especially salient. Beyond further work to account for population structure, `IBDGem` may also benefit from a more flexible model for sequencing errors, perhaps including the possibility of asymmetric errors or tetra-allelic (rather than biallelic) errors (Mostad et al., 2023).

Our examination of `IBDGem` has been limited to the statistical issues regarding the likelihood ratios discussed here. We have not considered aspects of `IBDGem` downstream of the likelihood ratio for identity, such as likelihood ratios for IBD1 or detection of IBD segments.

Some of our results use our own implementation of a haploid version of the `IBDGem` model, which we pursued for easier illustration. However, because the authors of `IBDGem` made their work open and available, we were able to explore our results in their software. This level of openness is admirable and not universal in forensic genetics (Chin et al., 2019; Edge & Matthews, 2022; Matthews et al., 2019).

We focused here on the values taken by the likelihood ratio when the hypothesis generally of interest to the prosecution—that a known person of interest is the source of the evidence—is true. It is also important to delimit the circumstances under which likelihood ratios computed by `IBDGem` could lead the evidence to appear to favor the prosecution hypothesis when the evidence should instead favor the defense hypothesis, or favor the prosecution hypothesis only very weakly. Exploration of this question should entail both mathematical approaches and realistic simulations of genetic variation (Baumdicker et al., 2021; Haller & Messer, 2019).

The tremendous advances in DNA extraction from challenging samples holds promise for forensic identification (Berglund et al., 2011; Kapp et al., 2021; Kayser & de Knijff, 2011; Vohr et al., 2015). At the same time, stakes in the courtroom are very high, and the statistical behavior of new methods needs careful validation to establish error rates as well as boundaries to their reliability. It is important that likelihood ratios used in case work are constructed with respect to clear hypotheses of forensic interest, and that they are well calibrated or, if in error, conservative.

## Code availability

Code used in this work can be found at https://github.com/fouerghi/IBDGem-LRs. We used `IBDGem` v2.0.2 available on GitHub to run all analyses.

## Acknowledgments

We thank N. Adams, G. Coop, V. Link, M. Przeworski, R. Rohlfs, N. Rosenberg, J. Schraiber, and members of the Edge, Mooney, and Pennell labs for helpful conversations. We acknowledge support from NIH grant R35GM137758 to MDE.

## Methods

### Diploid simulations using IBDGem

### Site-frequency spectra

To simulate haplotypes that adhere to a neutral site-frequency spectrum (SFS), we generated an allele-frequency distribution that matches the distribution expected under neutrality. We constructed this vector of allele frequencies by normalizing a vector of probabilities proportional to 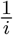 for *i ∈* {1, …, *n −* 1}, where *i* is the derived allele count in the sample.

To simulate a more realistic SFS, we used the 1000 Genomes Project VCF file for chromosome 1 available at https://ftp.1000genomes.ebi.ac.uk/vol1/ftp/data_collections/ 1000 genomes project/release/20190312 biallelic SNV and INDEL. We subsetted the original file into two separate files, one containing only the CEU population and the other containing only the YRI population. For each subpopulation, we subsequently filtered the VCF to only retain biallelic sites with a minor allele frequency of above 1%. We then extracted the allele frequencies for the remaining biallelic sites and divided them into frequency bins, excluding sites with allele frequencies of 0 or 1. We obtained a final vector of allele-frequency distributions by dividing the number of sites per bin by the total number of biallelic sites analyzed.

### Diploid genotypes

Building on the simplified haploid example, we simulated a diploid target individual by simulating two haplotypes that are paired to form two genomes. We used the allele-frequency distributions derived from either a neutral or more realistic human SFS to generate the two haplotypes. The target genotypes were kept constant across all figure panels except the panel varying the number of loci. Reference genotypes for Figure 2 were simulated in linkage equilibrium.

### Reads for diploid genotypes

In most panels of Figure 2, we generated exactly two reads per site, keeping the specific reads constant. In the panel exploring varied sequencing depth, the numbers of reads per site were drawn from a Poisson distribution parameterized by the sequencing depth. To obtain these reads, we used the genotypes of the individual of interest at variable sites and sampled from each allele randomly, picking the allele represented in the sequencing read uniformly at random if the site is heterozygous. We then drew sequencing reads from a biological source that matched the sampled allele at each locus with probability 1-*ϵ*; otherwise, reads did not match the sampled allele.

### IBDGem Inputs

We stored the genotypes of the individuals in the reference database in a `.hap` file required as input to `IBDGem`. We additionally generated the `.indv` file that has a list of the samples included in the reference database. `IBDGem` also requires a `.legend` file that has the IDs of the variable sites, their positions along the specific chromosome and the reference and alternate bases. To generate this file, we randomly sampled from the four nucleotides A,T,C and G for the reference base and then sampled from the remaining three nucleotides for the alternate base.

We provided the simulated reads for the target genotype in the pileup format. This requires the “base quals” column that indicates the probability of an incorrect base call in a specific position. We adjusted the values in this column to match the error rate parameter. We simulated the column “mapping quality” to be around 0, to indicate our assumption that the reads have not been mis-mapped.

### IBDGem Commands

To perform the `IBDGem` comparison between the DNA sequence and the individual of interest under linkage disequilibrium, we ran the command with the following options:

~~~
ibdgem --LD \
   --hap reference_panel.hap \
   --legend reference_panel.legend \
   --indv reference_panel.indv \
   --pileup unknown_dna.pileup \
   --pileup-name sample100 \
   --sample sample100 \
   --window-size 50 \
   --error-rate 0.02
~~~

We removed the –LD flag to run the comparison under linkage equilibrium. We adjusted the

–error-rate and –window-size flags depending on the parameters defined in the simulation.

#### Models including linkage disequilibrium

### Blocks of perfect linkage disequilibrium

#### In haploids

To simulate blocks of *m* loci in perfect linkage disequilibrium, we made *m−*1 copies of the originally simulated haplotypes. *P* (*D*|*I*) and *P* (*D*|*R*) for data consisting of one “0” read from each locus were computed as before, only on sets of *m ∗ k* loci. To compute the likelihood ratio assuming linkage equilibrium, we used the product rule to obtain *P* (*D*|*U*), including all loci. To compute the correct likelihood ratio accounting for linkage disequilibrium, we used

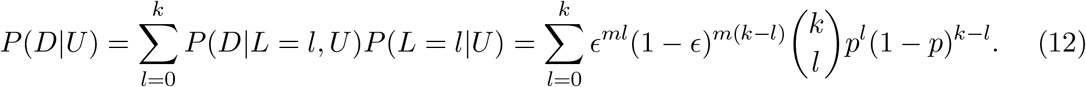

This reduces to equation 10 if *m* = 1. If *m >* 1, then in any blocks that do not match the individual of interest, there must be *m* sequencing errors to explain the data, leading to the *ϵ*^*ml*^(1 *− ϵ*)^*m*(*k−l*)^, where *l* is now the number of blocks, rather than the number of variable loci, at which an unrelated individual does not match the individual of interest.

#### In diploids

Similarly to the haploid model, we simulated blocks of *m* loci in perfect linkage disequilibrium by making *m −* 1 copies of the originally simulated genotypes. To compute the standard forensic likelihood ratio, we considered the reference database without the redundant markers which should represent loci in approximate linkage equilibrium and run `IBDGem` in non-LD mode, treating reads from sites in the same perfect-LD block as reads from a single site. To obtain the likelihood ratio assuming linkage equilibrium, we ran `IBDGem` in non-LD mode on the reference database with the sets of *mk* loci. Finally, running the `IBDGem` LD-mode on the reference database with the redundant markers resulted in `IBDGem`’s likelihood ratio computation.

### “Caterpillar” complete-linkage-disequilibrium haplotypes

#### In haploids

To simulate “caterpillar-like” complete-linkage-disequilibrium haplotypes, as exemplified in Table 2, we drew integers between 0 and *k* uniformly at random. The value drawn determines the number of “1” alleles at the start of the haplotype; the rest are assigned “0”.

*P* (*D*|*I*) and *P* (*D*|*R*) for data consisting of one “0” read from each locus were computed as before. To compute the likelihood ratio assuming linkage equilibrium, we used the product rule to obtain *P* (*D*|*U*), including all loci, and using the allele frequencies *k/*(*k* + 1), (*k −* 1)*/*(*k* + 1), …, 1*/*(*k* + 1). To compute the correct likelihood ratios, we used

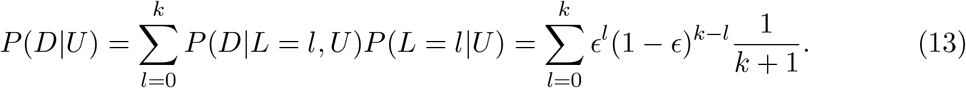

Here, *P* (*L* = *l*|*U*) = 1*/*(*k* + 1) because there are *k* + 1 equally frequent haplotypes, one with each possible value of *L*.

#### In diploids

Based on the number of loci in consideration, we simulated a reference database that includes all possible genotypes, each represented at their appropriate frequency. First, we simulated all possible “caterpillar-like” haplotypes, and then formed all possible genotypes by joining these haplotypes in every possible pair. We constructed the reference database with two copies of each heterozygous genotype to reflect that each heterozygous pairing has two possible orientations, and one of each homozygous genotype.

To obtain the standard forensic likelihood ratio, we ran `IBDGem` LD-mode on the entire reference database constructed as described above. We created a new reference dataset of size *n* by sampling with replacement from the entire reference dataset. We computed the likelihood ratio under the assumption of linkage equilibrium by running `IBDGem` non-LD mode on the downsampled dataset. We obtained the `IBDGem` simulated likelihood ratios by running `IBDGem` in LD mode on the downsampled dataset.

