## Supplementary figures for "On forensic likelihood ratios from low-coverage sequencing"

November 24, 2024

### S1 Supplementary Figures

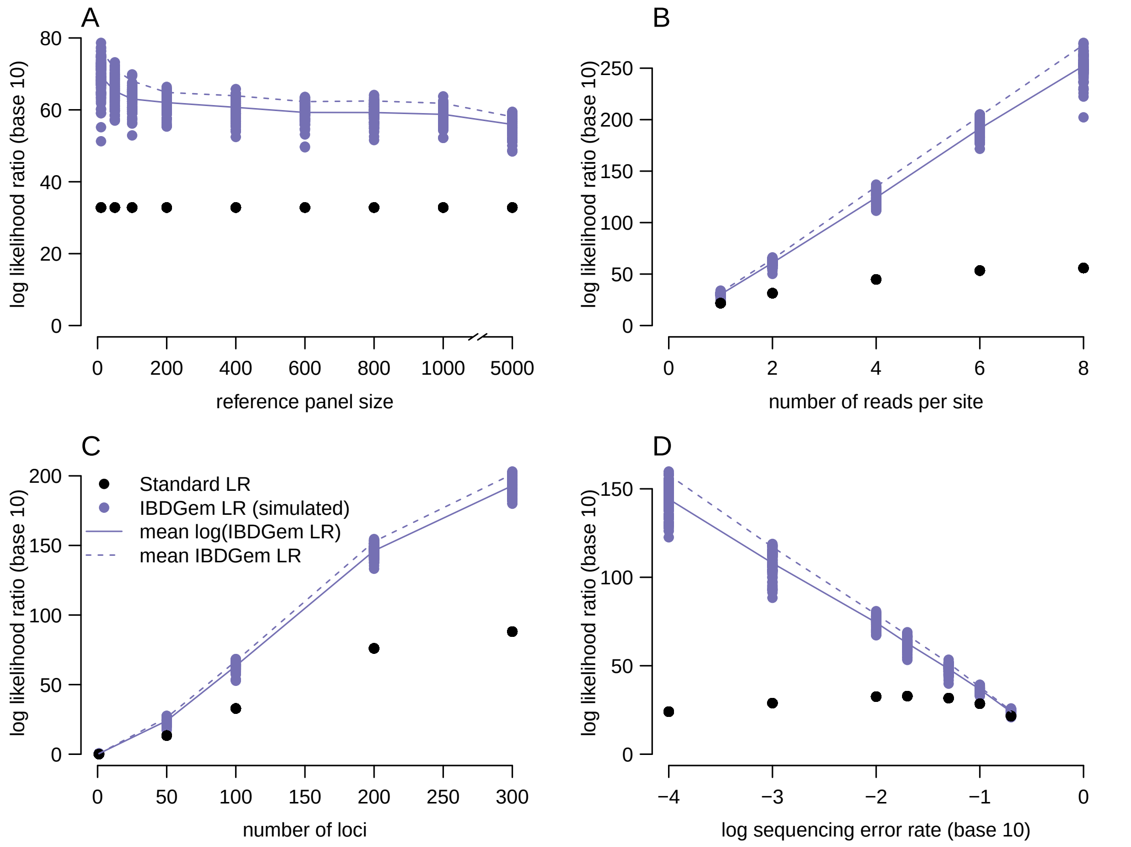


Figure S1: Dependence of the standard forensic likelihood ratio and `IBDGem` LD-mode likelihood ratio on parameters in a diploid setting with sites in linkage equilibrium obeying a site-frequency spectrum derived from the YRI subset of the 1000Genomes panel. These likelihood ratios were computed using `IBDGem` in either LD mode (purple) or non-LD mode (black). Conventions and baseline parameters are the same as in Figure 2 in the main text.

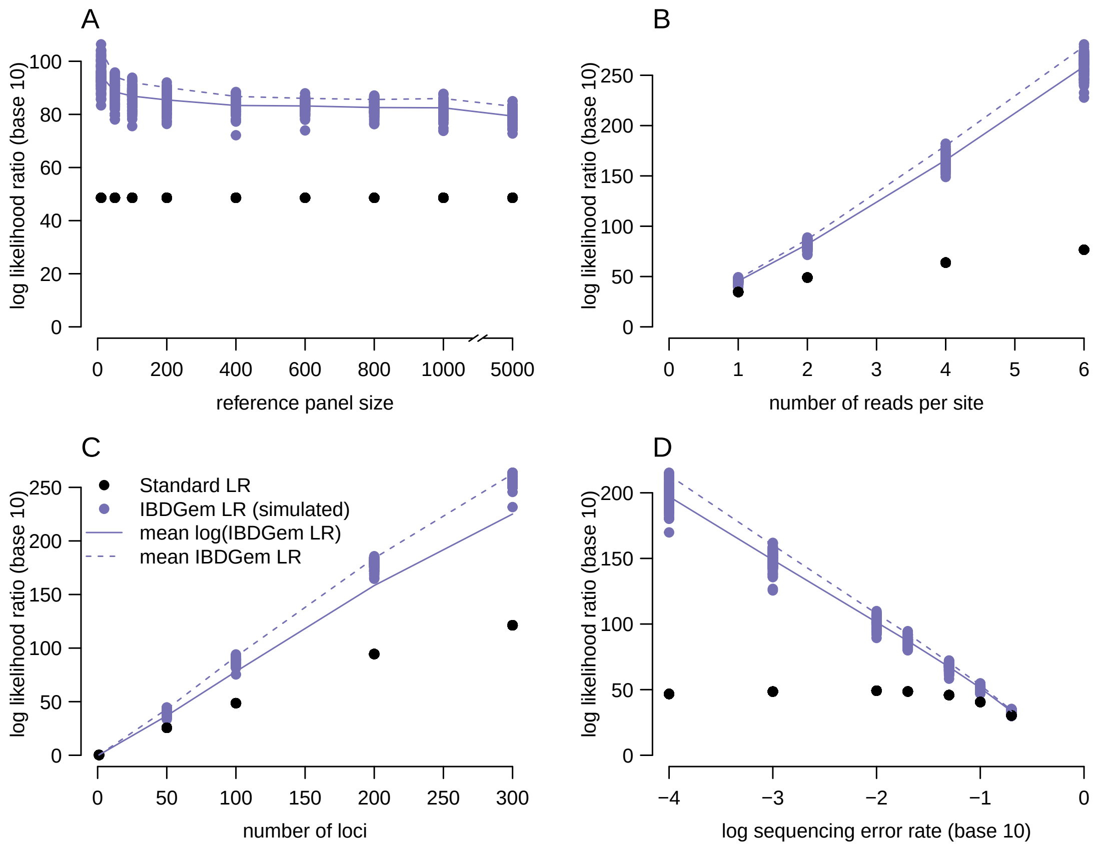


Figure S2: Dependence of the standard forensic likelihood ratio and `IBDGem` LD-mode likelihood ratio on parameters in a diploid setting with sites in linkage equilibrium obeying a site-frequency spectrum derived from the CEU subpopulation of the 1000Genomes panel. These likelihood ratios were computed using `IBDGem` in either LD mode (purple) or non-LD mode (black). Conventions and baseline parameters are the same as in Figure 2 in the main text.

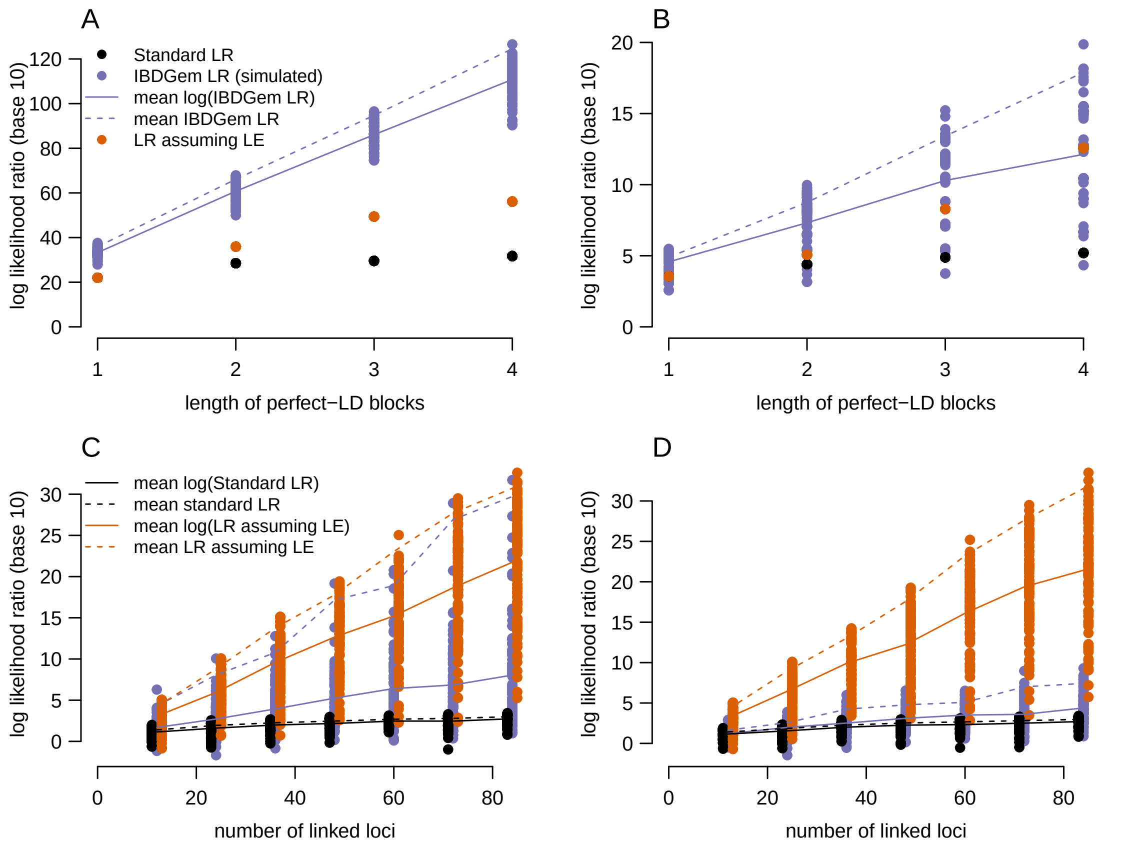


Figure S3: In the presence of linkage disequilibrium in a diploid setting, likelihood ratios following the model for `IBDGem`'s non-LD mode will overstate evidence for identity. Conventions are the same as in Figure 3 in the main text. Panels A and B show results from perfect blocks of LD, and panels C and D show results from the complete-LD model explained in the Methods section. In all panels, the sequencing error rate is $\epsilon = .02$, and there are exactly two reads per site. In panels A and B, the reference database size is $n = 100$. In Panel A, $k = 50$, whereas in Panel B, $k = 5$. In Panel C, $n = 20$, whereas in Panel D, $n = 100$. There are multiple dots for each of the likelihood ratios, because in the case of complete LD, the target genotype varies across simulations. (In the complete-LD model, results depend strongly on the specific genotype drawn as the target, so we drew multiple target genotypes for clarity.)

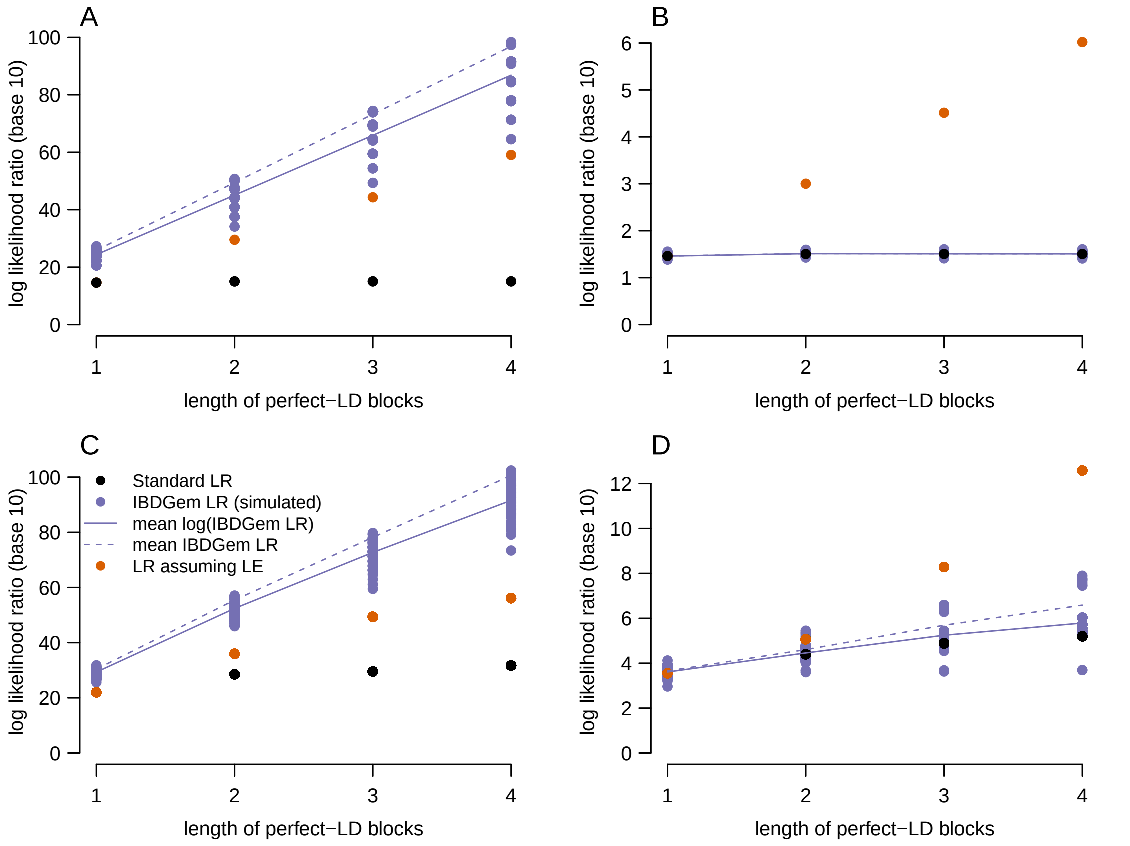


Figure S4: Comparing perfect-LD simulations in the haploid and diploid cases. Conventions are the same as in Figure S3. Panels A and B display results from perfect blocks of LD in the haploid setting, while panels C and D show diploid results. In panels A and C, $k = 50$, whereas in panels B and D, $k = 5$. In all panels, we set the size of the reference database $n = 5000$. The `IBDGem` LD-mode LR and the standard LR do not agree when using a window size of $k = 50$ in either the haploid or diploid simulations.

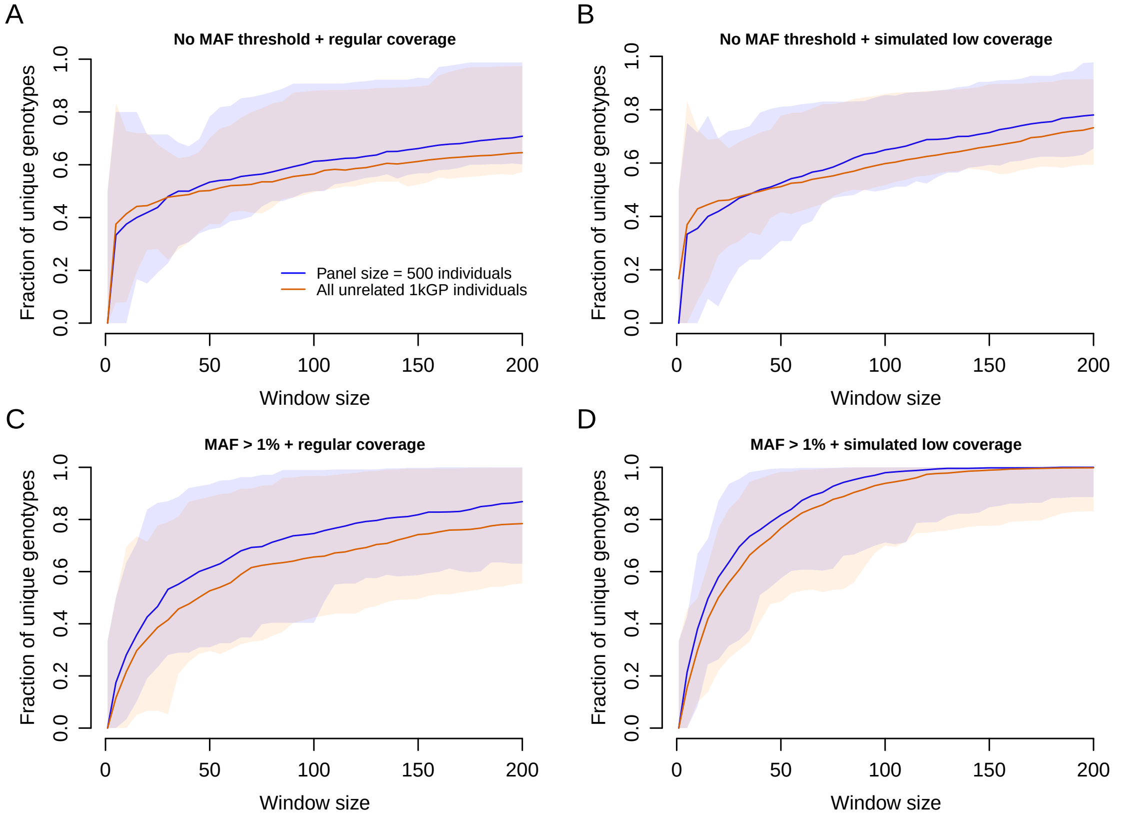


Figure S5: For moderately sized analysis windows, many or most multilocus genotypes in the 1000 Genomes project on chromosome 1 are unique. In all panels, the horizontal axis shows the window size (plotted points are windows of {1,5,10,15,20,...,200} variable sites), and the vertical axis gives a fraction of individuals with unique multilocus genotypes. In panels A and B, we retain all biallelic sites, while in panels C and D, we retain only sites with minor allele frequencies of at least 1%. Unlike panels A and C, where all biallelic sites on chromosome 1 are covered, panels B and D simulate low coverage by assigning each biallelic site a 10% probability of inclusion. We use two different panel sizes: one that includes all 2,504 unrelated individuals in the 1000 Genomes Project and another created by randomly subsampling 500 individuals. For each panel size, the line represents the median fraction of unique genotypes across 100 random windows, each initiated at a different starting location for the window on chromosome 1. The shaded regions show the minimum and maximum fractions of unique genotypes across those 100 random windows.

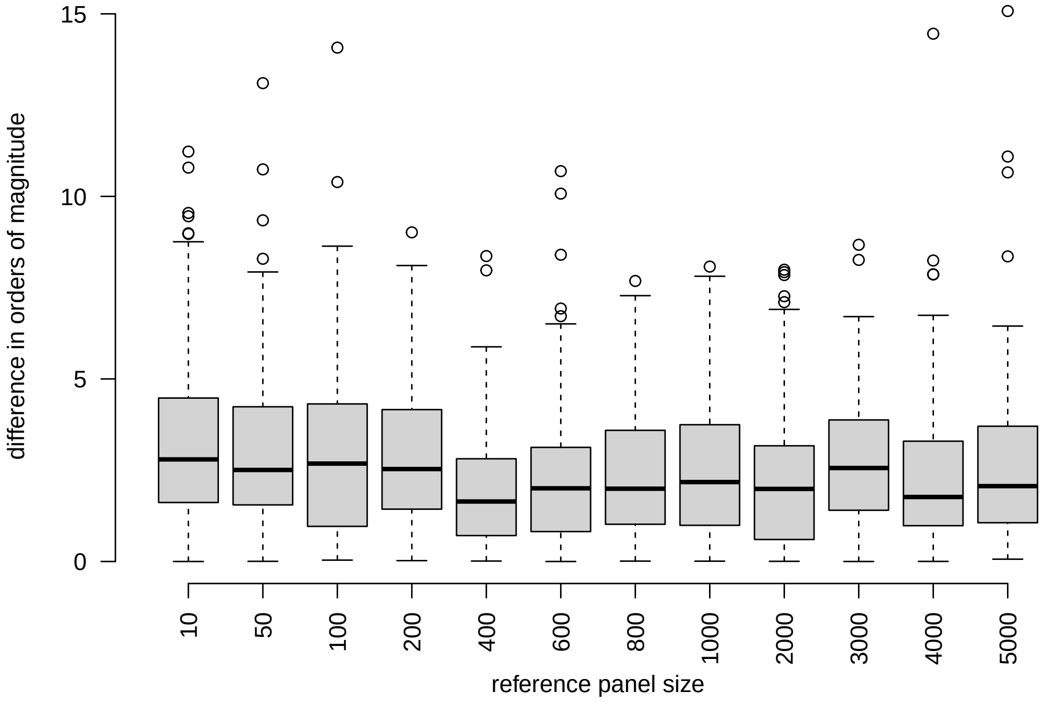


Figure S6: `IBDGem` is robust to phasing error (and indeed does not include phase information) but does not allow for any variation beyond the multilocus genotype frequencies observed in the reference database. Baseline parameters are number of loci $k = 100$, and sequencing error rate $\epsilon = 0.02$. This figure displays the order-of-magnitude difference between the LD-mode likelihood ratios calculated in the reference databases that result from reshuffling haplotypes within the panel to form new individuals.
